# Development of an integrated structural biology platform specialized for sub-100 kDa protein complexes to support biologics discovery and rational engineering

**DOI:** 10.1101/2021.03.24.436836

**Authors:** Yuri Iozzo, Yu Qiu, Albert Xu, Anna Park, Maria Wendt, Yanfeng Zhou

## Abstract

Developing a biologic medicine requires successful decision making at each step of selection, optimization, and/or combination of the right candidates at early research stages. Knowing the structural information and binding pattern between drug target and discovery candidates greatly increases the possibility of success. With the cryo-EM resolution revolution and rapid development of computational software, we have evaluated and integrated different tools in structural biology and the computation field and established a highly cost-effective platform, which allows us to obtain fast and accurate structural information for biologics projects with a close to 100% success rate and as fast as weeks turn-around time. Here we report four case studies selected from over 40 different protein structures and share how we integrate cryo-EM structure determination, computational structure modeling, and molecular dynamics simulation. With proper decision making and strategic planning, the platform allows us to obtain quality results within days to weeks, including sub-100 kDa complexes which are usually considered as a challenge due to their small size. Our utilization of this differential approach and use of multiple software packages, allows to manage priorities and resources to achieve goals quickly and efficiently. We demonstrate how to effectively overcome particle orientation bias by altering complex composition. In several of our examples, we use glycan density to facilitate interpretation of low-resolution 3D reconstruction and epitope mapping. Protein information plays an important role in our cryo-EM projects, especially in cases where we see significant challenges in obtaining high-resolution 3D maps.

## Introduction

Biologic drug molecules have gained a large market share and delivered various treatments to previously undruggable diseases. Due to the high molecular complexity and specificity of biologics, availability of a high-resolution structure in early research phase is crucial for in-depth understanding of the mechanism of action of a candidate, molecule design, mitigation of developability risks. Rapidly developing in silico maturation techniques also heavily rely on the structure availability [1-4].

X-ray crystallography has been and remains a staple structural biology tool in pharmaceutical research and rational drug design due to its ability to achieve atomic resolution and due to availability of light sources [5]. Particularly, thousands of structures of Fabs and heavy chain variable domain of a heavy chain antibody (V_HH_) in complexes with antigens are available in Protein Data Bank. However, unpredictable success rates in crystallography call into question if the method can be reliably used on its own for biologic drug discovery needs, which usually require high success rates and fast turn-around times. In addition, large amounts of well-designed protein fragments are required for crystallization screenings, leading to staggering costs and overextended timelines. Finally, X-ray structure does not guarantee its full biological relevance due to crystal contacts [6] and the presence of solvents and polymers.

The cryo-EM resolution revolution took off in 2013 with the advent of direct electron detectors and new data processing approaches [7]. Since then, the method quickly evolved and became a staple tool in drug discovery [8-10]. Advantages of cryo-EM include the elimination of lengthy crystallization attempts and the uncertainty of obtaining well-diffracting crystals. Cryo-EM yields a structure, free of crystal contacts, from molecules in near-physiological conditions. Although, the method is mostly successful in molecules with rather large molecular weight, cryo-EM has steadily demonstrated success in the sub-MDa range, eventually crossing the 100 kDa threshold [11, 12]. Many structures or structure ensembles, unsuccessfully sought after with X-ray crystallography, were solved by cryo-EM [13-17]. In biologics research, a broad range of complexes, such as antigens bound to Fabs or multiple V_HH_, are typically close to or heavier than 100 kDa and, thus, are suitable targets for cryo-EM [18].

In this article we discuss cryo-EM results of four soluble complexes (size range 65-200 kDa) and share our learnings from over forty antigen-antibody (or V_HH_) structures solved for various biologics project needs. Such complexes were assembled from soluble proteins or fragments of cell surface receptors and Fab(s) or V_HH_. We demonstrate how an integrated and optimized structural biology pipeline (Fig. 1, Table 1) can efficiently deliver structural information for sub-100 kDa complexes and advance projects, with close to 100% success rates and weeks of turn-around time. In addition, we also discuss common problems we have encountered.

**Table 1.** Summary of the case study.

| Project | Resolution, Å | Symmetry | MW*, complex (w/o glycans) | Glycan | Software pipelines | Application of structures |
| --- | --- | --- | --- | --- | --- | --- |
| Antigen A, Fab A | 3.6 | C3 (trimer) | 140-200 kDa | No | cisTEM | Developability engineering, MoA** elucidation |
| Antigen B, V<sub>H</sub>H B1 and B2 | 3.4 | C1 (monomer) | 80-90 kDa | Yes | cisTEM, Relion | Epitope-Paratope, MoA elucidation, multispecific design |
| Antigen C, Fab C | 4.8 | C1 (monomer) | 105-110 kDa | Yes | cisTEM | Epitope identification, MoA elucidation, affinity alteration |
| Antigen D, Fab D | 8-10 | C1 (monomer) | 60-70 kDa | Yes | cisTEM | Epitope identification, MoA elucidation |

\*MW – molecular weight
\*\*MoA – mode of action

**Figure 1.**
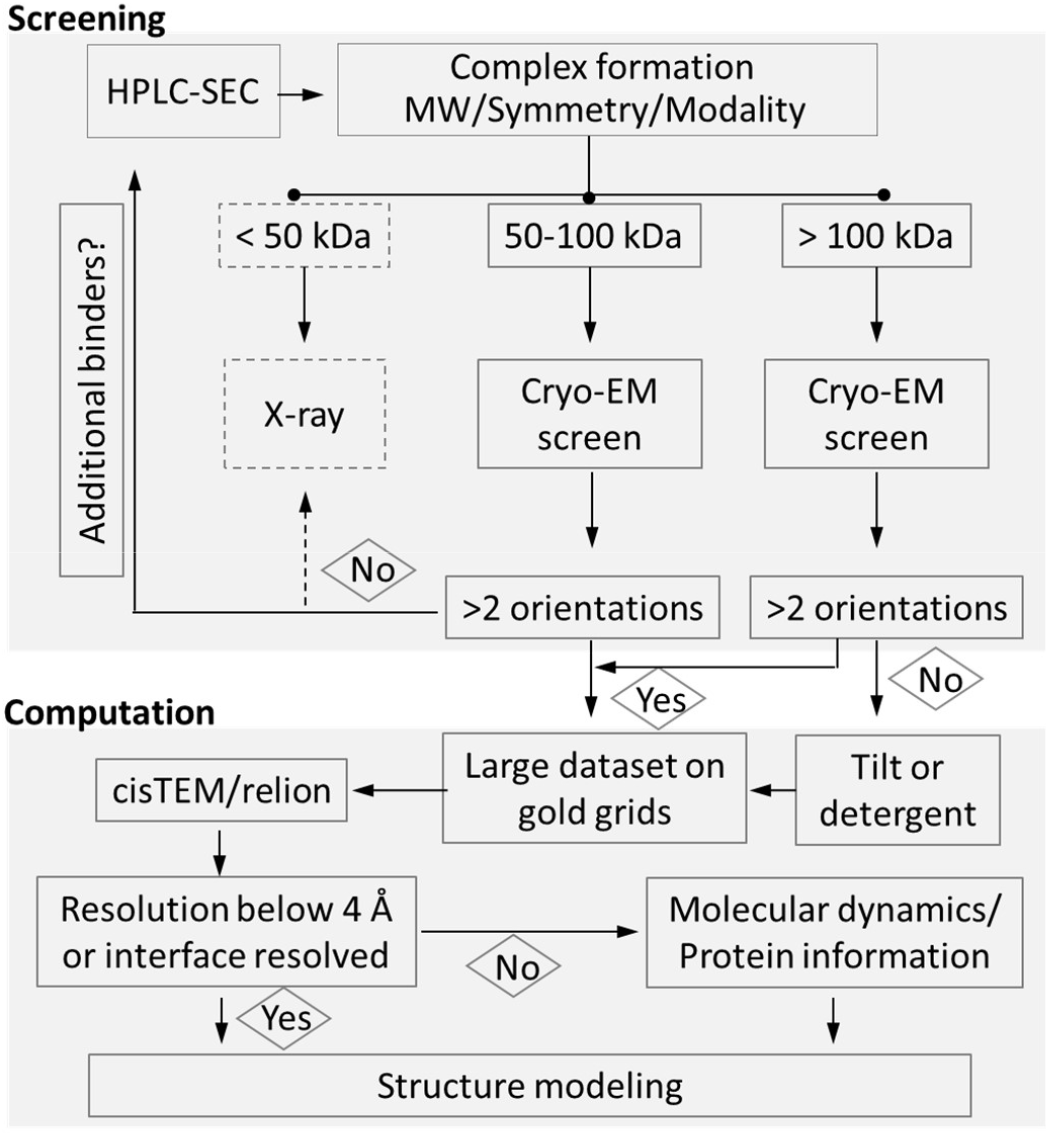
Flowchart showing the structural biology decision process. For complexes above 100 kDa, cryo-EM is the primary structure determination method. For smaller complexes, intrinsic antigen features like oligomerization and glycosylation are first considered during method selection. Symmetry greater than C1 usually leads to successful structure determination via single particle cryo-EM. If mitigation of strong preferred orientation for 50-100 kDa complexes fail, then crystallography is employed. Molecular dynamics simulation helps to resolve clashes and to relax the structure if cryo-EM resolution is lower than 4 Å.

## Results and Discussion

Requests for structural information of biologic candidates are diverse and, thus, can be filled flexibly in terms of deliverables. This contrasts with a mandatory need for atomic resolution for small molecule targets. Questions for biologic molecules include mode of action elucidation, binding orientation, epitope-paratope identification, affinity alteration, developability risk mitigation, etc. A promptly delivered medium-low resolution cryo-EM map, with the added support of computational structure modeling tools, may be sufficient to reveal candidate-antigen relative orientation and, often, to map the epitope-paratope interface. The following sections will focus on how we were able to deliver structural biology information of antigen-Fab/V_HH_ complexes, without a critical dependence on high resolution.

## 1. Optimized workflow for sub-100 kDa biologics complexes

### 1.1. Sample preparation

Use of commercial antigen materials saves a significant amount of resources. An increasing number of antigen proteins are available from different vendors, often, with fusion tags for bioassays. In contrast to crystallography, tags, such as flag or strep are well tolerated by cryo-EM. Obtaining an antigen from a commercial stock can save months of protein production efforts.

After a complex is formed and purified via size exclusion chromatography (SEC), sample grids are prepared from the purified complex using the Vitrobot system. Instead of going through time-consuming pH/buffer screening, a standard set of four screening buffers is used, which is suitable for most of our samples: each buffer contains NaCl and a buffering component (HEPES, Imidazole or Tris) with pH adjusted to 7.0, 7.5, 8.0 and 8.5 (see Methods). While these buffers may be sub-optimal for some samples, a limited number of screening buffers speeds up turnaround time to obtain the structure for most samples. Buffer or complex composition can be modified if severe preferred orientation is observed in 2D class averages (see more in sections 2.2., 2.4.).

Sample concentration is a critical factor to achieve an appropriate particle distribution on grids. Therefore, a range of concentrations is utilized in the first round of grid screening. It is worth noting that starting concentration of 0.1-0.2 mg/ml has proved to be a “sweet spot” for complexes and grids we use (UltrAuFoil, C-flat).

### 1.2. Grid screening without prior negative staining

The workflow is simplified by removing negative staining analysis of particles. Results from negative stained samples tend to fail to predict the particles’ optimization matrix under cryo-EM conditions or take as much time as screening and optimizing cryo-EM grids directly.

Two grid types, C-flat and UltrAuFoil, are preferred for water soluble biologics complexes. When a sample requires an extensive screening or the apparent size of a complex is large enough to produce a good signal to noise ratio, cost-effective C-flat grids are selected. However, when dealing with complexes smaller than 100 kDa, gold (UltrAuFoil) grids [19] are the first choice, and have proven to be most robust and reproducible in our pipeline, providing thin and stable ice. There are many other choices of grids, for example, coated with thin carbon or graphene oxide, however, these grids create high background noise or other challenges [20].

Mailing-out grids for screening and data acquisition at external providers greatly expedites the turn-around time. Because many target complexes fall into a size of around 100 kDa or below, only high-magnification capable Talos Arctica and Titan Krios microscopes equipped with a direct-electron camera are chosen for screening. This setup allows evaluation of 1-2 complexes within a single 24-hours instrument shift, including time for multiple grids screening and collection of two data sets.

### 1.3. Map generation

In fast-pace research, extracting sufficient information as quickly as possible is, often, more valuable than pushing the data to highest resolution. With on-the-fly motion correction and cloud data sharing, cryo-EM data is available after a data collection session as quickly as receiving X-ray diffraction data from a synchrotron.

Our workflow enables us to immediately start data processing with cisTEM [21] to quickly assess overall data quality, perform 2D classification, and generate an *ab initio* maps within a day or two. Two of the main determining factors to successfully obtain a final map via cisTEM are complex size and acceptable diversity of particle orientation. In many cases, cisTEM generates a reasonable quality map for model fitting, especially when diverse particle orientations are present (Fig. 2). At this point, 4.5-5 Å resolution is usually enough to predict paratope/epitope information and key residues on the antigen-Fab (or antigen-V_HH_) interface. Generally, local resolution at the interface is better than global resolution, as the interface is usually located near to the center of the mass of the complex. Once enough side chains are resolved to determine their conformation, further attempts to improve resolution cease.

**Figure 2.**
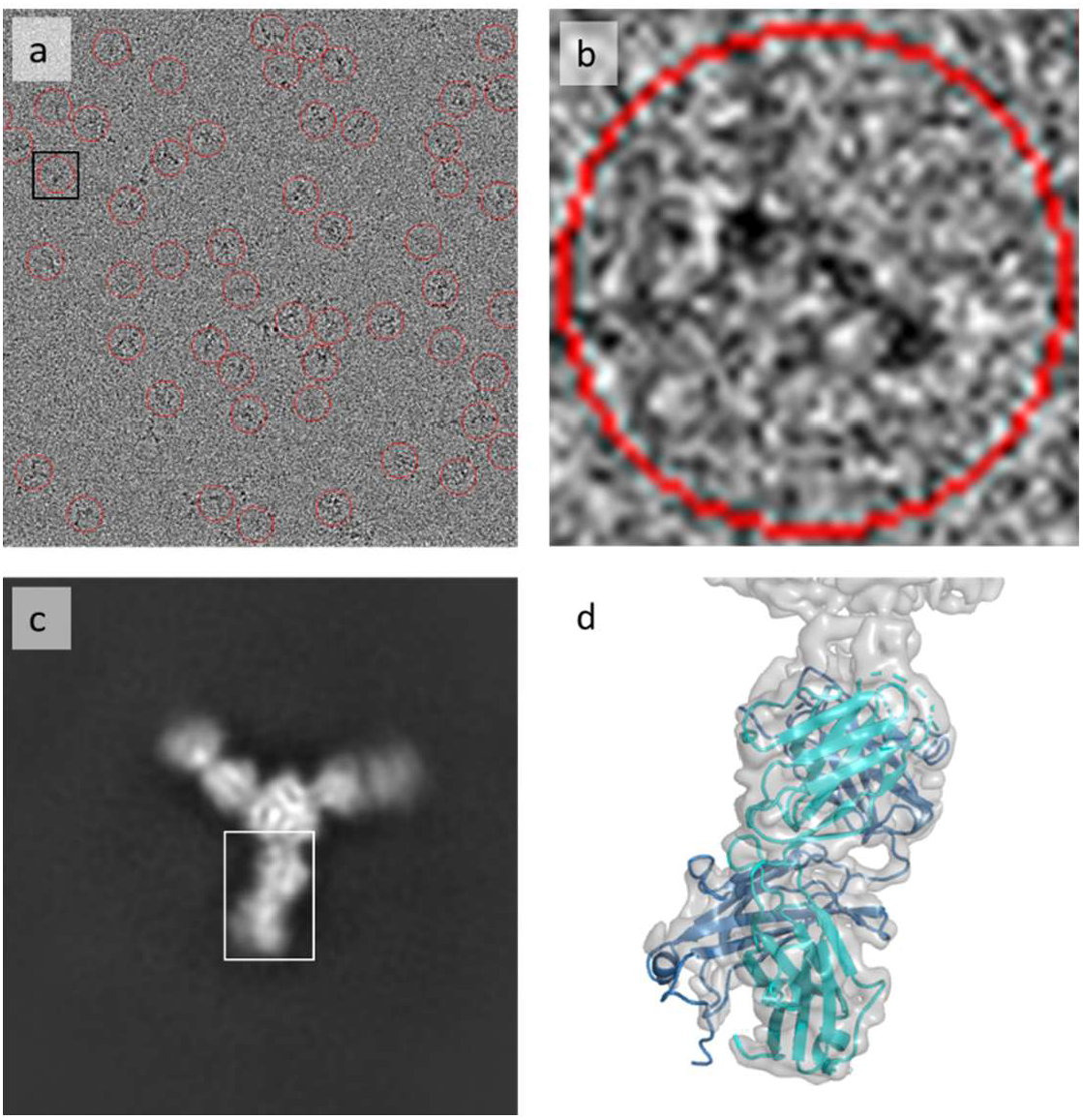
Structure determination of antigen A complex with Fab A solved at 3.5 Å. (a) A partial representative micrograph from Titan Krios microscope, equipped with K3 Summit camera. Particles picked by cisTEM are shown in red circles, the black square highlights the particle shown in panel b. (b) A particle showing C3 symmetry. The particle represents the “top” view of the molecule. (c) A class average (“top” view), showing C3 symmetry, obtained after 2D classification of particles in cisTEM. Fab and antigen are clearly distinguishable. The white box highlights the approximate area on the 2D image corresponding to the partial map shown in panel d. (d) A fragment of the reconstructed map at 3.5 Å. The Fab A model was refined in Refmac. Antigen is not shown. CDRs are removed.

Data processing will continue with Relion [22] if a higher resolution map is required. Relion jobs require significantly more time than cisTEM, so they may be fed with one or more cisTEM outputs: (i) images of 2D class averages can be used as a template input for particle picking via Gautomatch [23] with subsequent importing of coordinates to Relion, (ii) after 2D classification, a refinement package can be exported from cisTEM in STAR format, and particle coordinates from the package can be used for particle re-extraction in Relion, (iii) if cisTEM yields a well-resolved *ab initio*, such a map can be used as a reference for 3D classification or 3D refinement in Relion.

If we fail to obtain a 4-5 Å or better map in the Relion workflow, typically due to preferred orientation or complex size, we return to a sample optimization step to optimize conditions on the grid or to change sample composition by adding new components, as described in section 2.2.

### 1.4. Model building

Model building starts from rigid body fit of antigen, Fab, or V_HH_ models in a map using Chimera [24]. For this purpose, Fab and V_HH_ models are generated using MOE [25] Antibody Modeler. Framework regions of Fab and V_HH_ models are mostly reliable due to strong structural conservation. Detailed conformations of CDRs can be determined later once the models are fit into a map based on framework regions. If antigen structure is not available from PDB, it can be generated through homology modeling. Flexible antigen models are split into smaller domains to ensure a robust rigid body fit into a map.

Two methods of model refinement have been developed, with their use dependent on the map resolution. For high-resolution maps (better than ∼4 Å), rigid body fitted structures are further improved in Coot [26] via manual or automatic fit of CDRs and out-of-density regions. Subsequently, models are refined in Refmac [27] within CCPEM package [28]. For lower resolution maps (4-5 Å), final models are derived from rigid body fitted structures through molecular dynamics simulation in MDFF [29] and refinement in Refmac.

## 2. Representative Case studies

### 2.1. Antigen A with Fab A at 3.5 Å (trimer, C3 symmetry)

The first case study is a complex of Fab with a 15-20 kDa commercially available non-glycosylated antigen. The physiological trimer formation of antigen greatly increased the feasibility of utilizing cryo-EM on this target. In contrast to the tendency of preferred orientation in trimeric HA-Fab complexes [30], this complex showed diverse particle orientation on grids (Fig. 2a), although the overall shape of the complex is flat. The 3-fold symmetry (Fig. 2b, Fig. 2c) and various orientations allowed fast and straightforward 3D map generation (Fig. 2d) within a day of computation from a single data set.

### 2.2. Achievement of 3.4 Å resolution for 80-90 kDa complex (C1 symmetry)

Here we present a strategy for antigens that are small and do not have intrinsic oligomerization features. Many biologics targets are monomeric proteins, and those targets commonly give severe (only one dominant view in 2D class averages) or mild (2 or 3 dominant views) preferred orientation. Any skewed reconstruction due to preferred orientation will likely lead to clash of coordinates around the antigen-Fab/ V_HH_ interface during model building and raise challenges when identifying and interpreting the epitope-paratope interactions.

A high-resolution structure was needed for target B with V_HH_ B1. The monomeric antigen is 60-65 kDa and heavily glycosylated, resulting 10-15 kDa attributed to attached glycans. V_HH_ B1 is roughly 13 kDa.

A complex of antigen B with V_HH_ B1 was placed on C-flat grids and data was collected using a Titan Krios microscope. Very few 2D orientations were observed; however, high resolution features were clearly resolved, including easily distinguishable glycans and orientations of beta sheets. Multiple attempts to generate a meaningful *ab initio* reconstruction failed. To overcome the preferred orientation problem, the same grid was again used to obtain another set of movies collected with 30° stage tilt and without it. 2D classification of “tilted” images presented fewer high-resolution details in 2D classes and did not provide large classes with additional views. The combined dataset only led to a 5 Å reconstruction map that appeared streaky and distorted in one dimension. As a result, the rigid body fitted antigen and V_HH_ clashed so much around the interface that only the relative orientation of V_HH_ could be revealed. Exact epitope information as well as details of antigen-V_HH_ interface remained elusive.

From the same V_HH_ discovery campaign a second V_HH_, B2, was characterized as a different epitope binder. We reasoned that addition V_HH_ B2 (MW roughly 18 kDa, including 5 kDa attributed to the flexible C-terminal tag) to the complex would increase molecular weight and change properties of the complex to overcome the preferred orientation issue. cisTEM 2D classification of the data set collected from UltrAuFoil grids on a Titan Krios revealed classes showing high-resolution features of the complex. We observed 2D class averages corresponding to a view (Fig. 3a), closely resembling the dominant views from the single V_HH_ complex. However, many class averages corresponded to new, more rare views (Fig. 3b). Despite that, only particles selected after multiple rounds of extensive 2D classification yielded a meaningful *ab initio* map. After refinement in cisTEM we have obtained the final 4 Å map with many helixes, β-sheets, and side chains readily distinguishable. Models of both B1 and B2 V_HH_ were placed and oriented confidently during the rigid body fitting. Despite initial promise, the map demonstrated lower resolution in epitope areas and details of antigen-V_HH_ interfaces remained enigmatic. Homology modeling of CDR H3 for B2 V_HH_ was not producing a conformation reminiscent of observed density, and the resolution of the map precluded manual placing of residues. The final cisTEM map did not address project needs. Due to the presence of high-resolution features in diverse 2D classes, we decided to spend time and resources to re-process the whole data set in Relion 3.0. The best twelve 2D class averages from cisTEM, representing diverse views, were used as a template to pick particles via Gautomatch. After one round of 2D classification in Relion, classes were selected very loosely, removing only obvious junk classes. Several rounds of 2D and 3D classification were performed subsequently, with classes being selected more conservatively. Generation of *ab initio* and subsequent 3D refinement yielded a 3.6 Å map. Still noisy, the map was clearly superior to 4 Å map from cisTEM. Bayesian polishing allowed an increase in resolution to 3.4 Å, and deepEMenhancer [31] processing of half-maps reduced most of the noise and provided a very good B-factor sharpening (Fig. 3c). The final map demonstrated density for a bulk of the side chains (Fig. 3d) and provided clear information about antigen and V_HH_ conformations, epitopes, and CDR H3 of V_HH_ B2.

**Figure 3.**
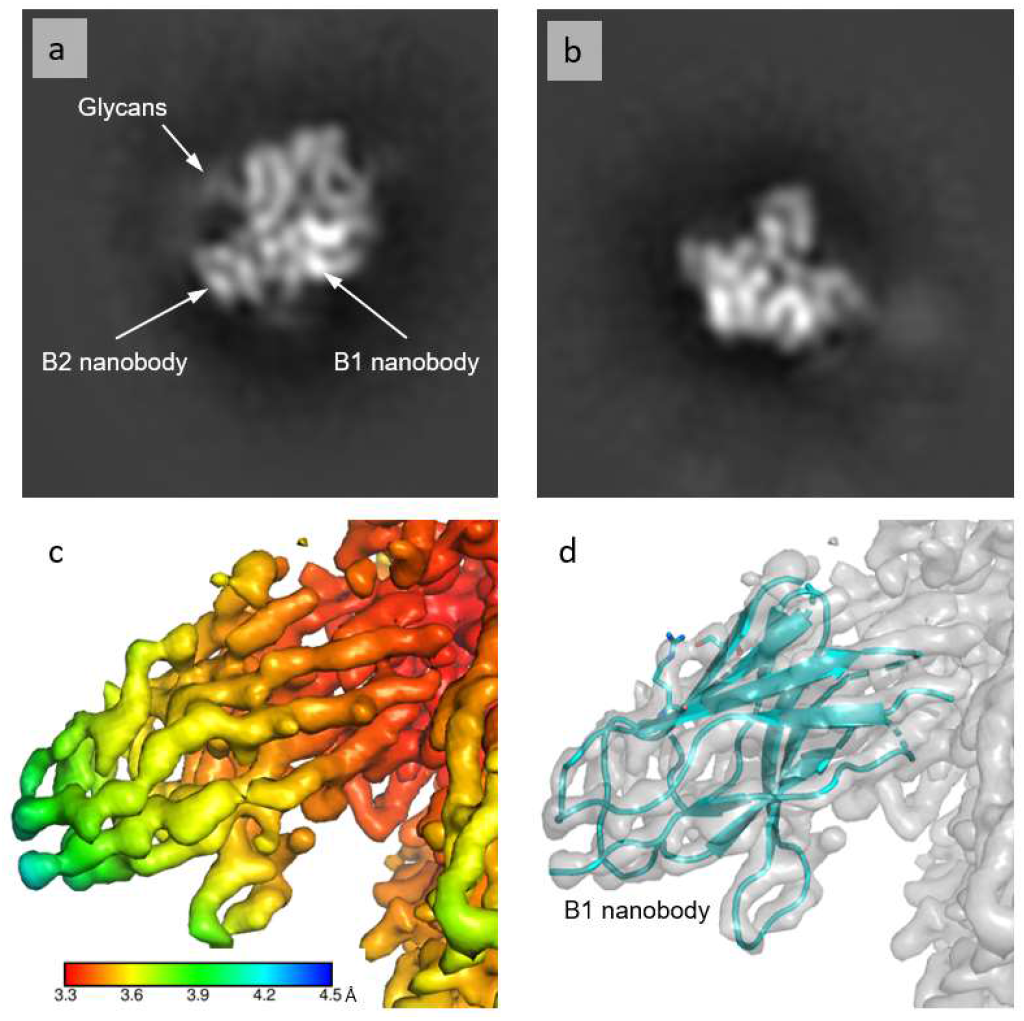
Structure of antigen B bound to two V_HH_ at 3.4 Å. (a) A 2D class average (horizontally flipped) representing a dominant view of the antigen B complex with two V_HH_, B1 and B2, obtained after 2D classification of particles in cisTEM. Low resolution features, including V_HH_ and glycans, are clearly visible. (b) A 2D class average (horizontally flipped) representing a rare view of the antigen B complex with two V_HH_, B1 and B2. (c) A fragment of the reconstructed map at 3.4 Å colored according to the local resolution. (d) Same map showing the model of V_HH_ B1 with two residues shown as sticks to highlight the resolution. CDRs are removed from the model.

Searching through EMDB entries showed that, as of December 2020, this map sets a record of high resolution for a structure with C1 symmetry with a MW at or below 90 kDa.

### 2.3. Antigen C complex with Fab C at 4.8 Å resolution (C1 symmetry)

In this case study, the complex was reconstituted from commercially available antigen C bound with Fab C and was put on C-flat grids. Two factors were screened sequentially: (i) buffer pH and composition and (ii) concentration of the complex. It appears that using imidazole buffer with pH 7.5 was critical for this sample, as all three other buffers produced aggregation on the grids. Complex concentration from 0.15 to 0.6 mg/ml did not dramatically affect the particle distribution on the grids; however, a higher (0.4-0.6 mg/ml) concentration was used for data collection because it seemed to produce more particles per micrograph (Fig. 4a). Notably, both thin and medium thickness area were checked during screening to find optimal area for data collection.

**Figure 4.**
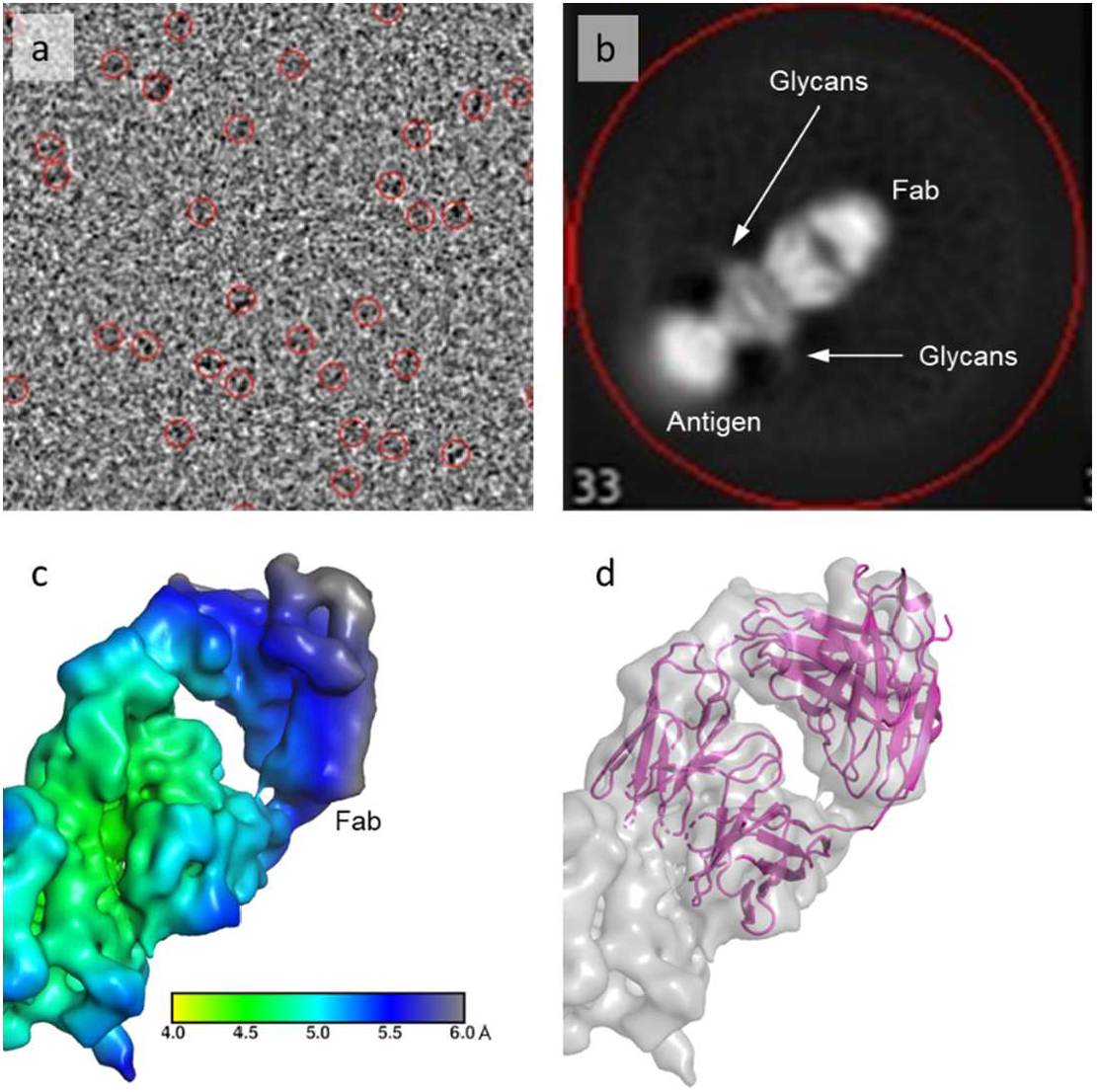
Structure of antigen C bound to Fab C at 4.8 Å. (a) A partial representative micrograph from a Titan Krios, equipped with K3 Summit camera, with particles picked in cisTEM (red circles). (b) A 2D class average showing Fab C bound to the antigen C. The distinct shape of the Fab as well as antigen glycans are clearly recognizable. (c) A fragment of the reconstructed map at 4.8 Å colored according to the local resolution. (d) The same map with Fab model refined into it in Refmac. Antigen is not shown. CDRs are removed.

A small dataset containing ∼750 movies was recorded from a Titan Krios and processed in cisTEM. After two rounds of 2D classification, high-resolution features became clearly distinguishable (Fig. 4b). However, particles demonstrated an orientation bias. Collecting an additional ∼1,600 movies with a 30° stage tilt did not improve the results. A low-resolution map was obtained from the merged data sets, which did not address the project needs.

A larger conventional dataset of ∼4,500 movies was collected in hopes of enriching 2D classification with rare views. After 2 rounds of 2D classification, remaining particles showed multiple orientations. Particles were exported to Relion 3.0 for 3D classification. A final 4.8 Å map was obtained after 3D refinement and post processing (Fig. 4c). It was sufficient to support model building and identification of epitope-paratope interactions. Due to its lower resolution, the final model was relaxed in MDFF before refinement in Refmac (Fig. 4d). The best data for this complex was always obtained from freshly prepared complex, thawing of a previously frozen sample for grid preparation led to low quality screening data.

### 2.4. Antigen D (20 kDa) complex with Fab D (C1 symmetry)

This is one of the smallest monomeric complexes we have encountered. Molecular weight of the antigen D complex with Fab D is roughly 65 kDa. With foreseen difficulties, we tried cryo-EM for several reasons: (i) a non-glycosylated complex failed to produce crystals under a tight project timeline; (ii) insights about glycans of antigen D and their effect on the antibody binding were needed. Many approaches were attempted to overcome issues throughout the analysis and although only a low-resolution map was obtained at the end, the resulting map provided valuable information.

The cryo-EM work was started utilizing non-glycosylated antigen from a prior crystallography study. The non-glycosylated complex (0.15-0.3 mg/ml) was applied onto R2/2 C-flat grids. Like in previous cases, various pH, buffers, and protein concentrations were screened. Grid screening demonstrated that particle distribution is more uniform at pH 8.0-8.5, and protein concentration did not severely affect particle distribution. A dataset containing about ∼3,800 movies was collected right after the screening. Over 1.4 million particles were picked and sorted by multiple rounds of 2D classification by cisTEM. However, only one dominant particle orientation was revealed in the 2D class averages. Although there were some slightly rotated views available, 3D reconstruction was not successful.

A few commonly used strategies were then applied to improve the particle orientation: (i) changing sample composition from non-glycosylated to glycosylated antigen, (ii) adding detergent, (iii) stage tilting during data collection, (iv) changing sample buffer.

In solution, glycosylated proteins may behave differently compared to their non-glycosylated counterparts [32]. Glycans can also provide additional landmarks for structural modeling. A batch of grids with glycosylated antigen D was prepared utilizing UltrAuFoil grids. More than two million particles were picked from ∼3,300 aligned movies. Although all three glycans were clearly seen, 2D classification revealed the same dominant views as observed in the non-glycosylated complex (Fig. 5a).

**Figure 5.**
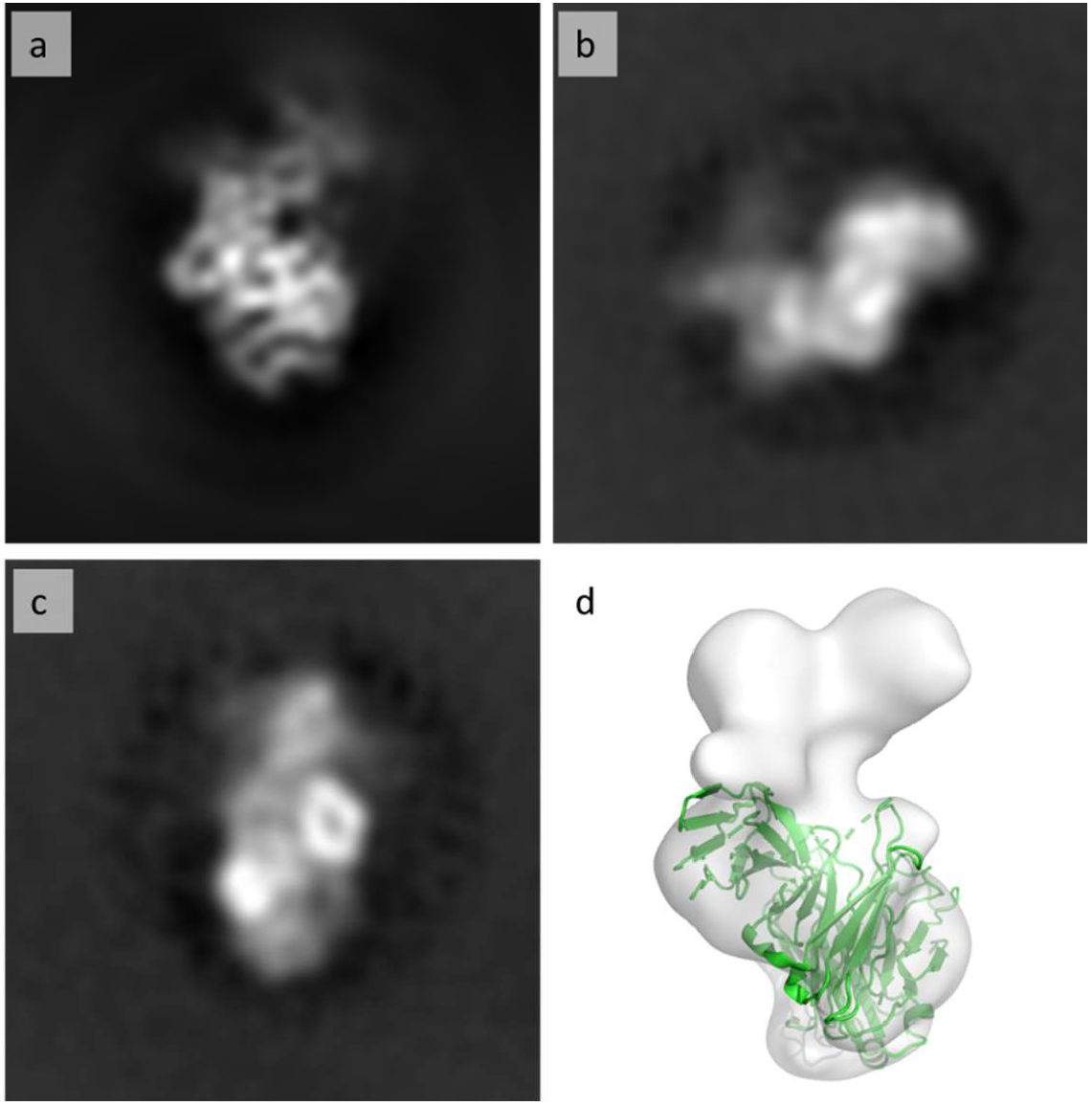
Structure elucidation of antigen D bound to Fab D (>6 Å). (a) A dominant particle view obtained after 2D classification of particles in cisTEM, present in all data sets. (b) A less-dominant particle view from a data set, collected in the presence detergent. (c) A less-dominant particle view from a data set, collected with a 30° stage tilt. (d) Final low-resolution map (>6 Å) with a Fab model, relaxed via MDFF. Antigen is not shown. CDRs are removed.

A few different orientations were obtained on UltrAuFoil grids after introducing detergents or stage tilting. With 30-40° stage tilting, more orientations emerged and, interestingly, represented mainly side views of the Fab (Fig. 5b). Addition of detergent, even at low concentration, produced a key orientation: viewing through the Fab doughnut hole (Fig. 5c). These additional views were necessary to calculate 3D reconstruction at low resolution (Fig. 5d).

We have used a publicly available structure of antigen D from Protein Data Bank and a MOE-generated Fab model to coarsely assign the epitope on antigen D guided by the glycans. The proposed model was improved by fitting into the low-resolution 3D volume. It has provided sufficient epitope information to advance this project. Therefore, the strategy of adding additional binders, described in section 2.2., was not explored in this case.

In this case, all attempts to obtain a high-resolution reconstruction failed, and we suspect that the signal to noise ratio in tilted or detergent data was significantly compromised and aggravated by the small size of this non-symmetric molecule. With lessons learned on this target, we find that for our sub-100 kDa complexes, once preferred orientation occurs, it is critical and effective to introduce additional binders to the target first before spending time on exploring detergent or collecting data with a tilted stage.

## 3. Methods

### 3.1. Sample preparation

Antigens, antibodies, and V_HH_ were purchased from commercial sources or purified internally. Antibodies were digested with FabULOUS or FabRICATOR enzymes (Genovis). Complexes were assembled in PBS at room temperature by incubating antigen with 1.2-1.5 molar excess of Fab or V_HH_ for 45-60 minutes. In case of antigen B, the second V_HH_, B2, was added 45 minutes into the initial incubation, the resulting mixture was incubated for an additional 45 minutes at room temperature. Complexes were immediately purified through SEC (Superdex 200 Increase 10/300 GL, Cytiva) on an Akta Basic (Cytiva) via isocratic elution in PBS using a 0.5 ml injection loop. Fractions, 0.25 ml each, were collected into 96 deep-well blocks. Fractions of the peak with the lowest retention time corresponded to a complex and were pooled together. Complexes were concentrated on Amicon 10-50 kDa cutoff 5 ml centrifugal filters (Millipore) to a volume of 100— 250 μl. In some cases, the volume was further reduced with an Amicon 10 kDa cutoff 0.5 ml filters (Millipore) down to 50 μl. Complex concentration was measured via Nanodrop (Thermo Fisher) and varied in the range of 0.5-8.0 mg/ml. To make grids for initial screening, samples were diluted by one of the following four buffers: 20 mM HEPES, 150 mM NaCl pH 7.0; 20 mM imidazole, 150 mM NaCl pH 7.5; 20 mM Tris-HCl, 150 mM NaCl pH 8.0; 20 mM Tris-HCl, 150 mM NaCl pH 8.5. If the concentration of the complex in PBS was too low to dilute it at least 7-10 times with a buffer before grid making, then Amicon 10 kDa cutoff 0.5 ml centrifugal filters were used for buffer exchanging. Samples with a final concentration of 0.1-1.0 mg/ml were centrifuged for one minute at 14,000g. If a detergent was used, it was applied right before grid making by adding 20x detergent solution to the sample, mixing and centrifuging again. UltrAuFoil R0.6/1 and R1.2/1.3 300 mesh (Quantifoil Micro Tools) and C-flat-2/2-2Au (Protochips) grids were used for grid making. Grids were glow-discharged at −20 or −25 mA for one minute using a PELCO easiGlow glow discharger (Ted Pella). 3-4 μl of samples were applied onto grids using a Vitrobot (Thermo Fisher). Grids were blotted with “595” filter papers (Ted Pella) for 4-9 seconds with blot force 1-10 (higher blotting time and force for C-flat grids, lower for UltrAuFoil grids) at 85-100% humidity and then plunged into liquid ethane. Specific details for each complex are shown in the following paragraphs.

The complex with antigen A was formed by incubating the antigen with 1.5 molar excess of Fab for an hour at room temperature. It was purified via SEC and applied onto UltrAUfoil and C-Flat grids in different buffers with pH ranging from 7.0 to 8.5, as described above. The complex concentration on the grids varied in the range 0.1-1.0 mg/ml.

The complex of antigen B with the single V_HH_, B1, after SEC purification, was applied onto C-flat grids at concentrations 0.2-0.4 mg/ml in all four buffers mentioned above. The complex with two V_HH_ was prepared from the antigen and 1.5 molar excess of each V_HH_ (B2 was added 45 minutes after B1) and purified via SEC. The complex was diluted in four testing buffers to 0.15-0.8 mg/ml and then applied onto UltrAuFoil grids.

The complex with antigen C, was prepared with 1.5 molar excess of Fab C. After SEC purification, the complex was applied onto C-flat grids in four buffers (listed above) with the concentration ranging 0.15-0.6 mg/ml.

The complex of antigen D, prepared in a two-fold molar excess of Fab D, was purified via SEC and concentrated to ∼8 mg/ml. It was then diluted in buffers, as described above, to 0.15-0.3 mg/ml and used to prepare UltrAuFoil grids for screening and collection of the first data set. The second set of C-flat grids was prepared using same buffer (20 mM Tris, 150 mM NaCl pH 8.0) and complex concentration of 0.2 mg/ml for each grid, but with the following final concentrations of detergents added: 0.2 or 0.5 mM CTAB, 1 or 3 mM fluorinated fos-choline-8, 1 or 2.4 mM ß-octylglucoside, 1 or 3 mM CHAPS.

### 3.2. Grids screening and data acquisition

We have access to Talos Arctica and Titan Krios microscopes (Thermo Fisher) equipped with K3 Summit cameras (Gatan) and SerialEM software [33]. The Talos Arctica microscope is mainly used for grid screening. For most complexes, grid screening and data collection were performed on the Titan Krios within one microscope session. Data acquisition is performed via multi-hole technique for R1.2/1.3 or smaller grids, or multi-hole and multi-shot approach for grids R2/2 or larger [17].

For the complex with antigen A and Fab A, a single data set of 3,771 movies was collected.

For the complex of antigen B with the single V_HH_, B1, three data sets were collected from the C-flat grid with a complex concentration 0.2 mg/ml: 642 movies collected using a Volta phase plate [34], 1,911 movies collected using conventional technique and 1,847 movies collected with and without 30° stage tilt (mixed data set). For the complex of antigen B with two V_HH_, a single data set was collected from the UltrAuFoil grid.

For the complex of antigen C with Fab C, three data sets were collected: a smaller data set of ∼750 movies, a data set containing ∼2,350 movies collected with and without 30° stage tilt, and a large conventional data set with ∼4,500 movies.

About 3,800 movies were collected for non-glycosylated antigen D bound to Fab D. A set of ∼3,300 movies was collected for the glycosylated complex. Later, more data sets were collected with a tilted stage and from the sample with detergent.

### 3.3. Data processing

Our cryo-EM partners provide motion correction on the fly through Relion or IMOD [35]. For early projects, we have directly imported IMOD-corrected movies for cisTEM processing. However, there are few problems (missing metadata, different motion-correction protocol) if Relion processing is required down the road. To avoid this, we now import only Relion motion-corrected images to cisTEM. Within cisTEM, the CTF estimation was done via default options. Particle picking required some optimization of maximum and characteristic particles radii. For all complexes, particles were picked at 15 Å resolution because using higher resolution for sub-200 kDa complexes often results in incorrectly picked particles. 2D classification, *ab initio* map generation, automatic refinement, and B-factor sharpening were done predominantly with default parameters (the hand was inverted, if needed, during the sharpening).

Data processing in Relion 3.0 was performed exactly as described in the software tutorial with the exception of particle picking. We run Relion on 8 GPUs and 30 threads. Particle positions were imported from cisTEM (for antigen C) or picked via Gautomatch v0.56 (for antigen B) using 12 different templates showing different molecule projections, obtained from cisTEM-generated mrc files containing images of 2D class averages. These templates were extracted from mrc files using v2 and proc2d scripts from EMAN/EMAN2 [36]. If needed, final volumes were inverted through relion_image_handler and the ‘--invert_hand’ option. More details for each complex are provided below.

For antigen A and Fab A, after quick visual inspection, 1,019 motion-correct images were removed due to excessive drift, icing, or unacceptable FFT. In hindsight, this step appears to be unnecessary. Resulting MRC images were imported to cisTEM, CTF was determined, and particles were picked. Several rounds of 2D classification led to selection of the best classes showing many expected structural features of the complex (Fig. 2c). Both *ab initio* reconstruction and Auto 3D refinement were performed in C1 symmetry, yielding a 4.1 Å map. Subsequently, the map was refined again but with C3 symmetry, which improved the resolution to 3.5 Å.

For the antigen B complex with B1 V_HH_, the Volta phase plate failed to produce quality results and was discarded. The second conventional data set demonstrated preferred orientation, which was not dramatically resolved by using 30° stage tilt in the third data set. *Ab initio* was obtained from the third set, and the final map demonstrated 5 Å resolution. For antigen B complex with two V_HH_, 2D class averages were obtained in cisTEM and used as Gautomatch templates. Further processing was done in Relion. The final 3.4 Å map (before the postprocessing stage) was processed via deepEMenhancer from two half-maps using the highRes option to sharpen B-factor and remove noise.

For antigen C, all three data sets were processed in cisTEM. The first two were eventually discarded. For the third and largest data set of ∼4,500 movies, two rounds of 2D classification were performed in cisTEM, after which ∼420,000 particles were exported to Relion 3.0 for 3D classification. The final 4.8 Å map was reconstructed after 3D refinement and post processing from ∼240,000 particles (Fig. 4c).

For antigen D, high quality 2D class averages (Fig, 5a) and a meaningful low-resolution map (Fig. 5b) were obtained from the second data set (the conventionally collected data for the glycosylated complex). The rest of the data was discarded because it failed to provide any improvement.

### 3.4. Model building and refinement

For all complexes described in this manuscript, antigen models were obtained from Protein Data Bank. For projects not described in this manuscript, where such models were not available, we have obtained a homology model in MOE using structures closest known homologs. All Fab and V_HH_ models were generated in MOE Antibody Modeler. All models were fit as rigid bodies into their corresponding maps in Chimera. CDRs and out-of-density antigen loops were manually fit into maps in Coot. For antigens A, B, and C, final models were refined in Refmac within the CCPEM package. For antigen D, the rigid body fit was obtained using Molrep [37] from the CCPEM package. Glycans and other structural features of the complex, clearly visible on 2D class averages, were used as landmarks via a proprietary method. For the complex with antigen C, molecular dynamics simulations for 50 ns in MDFF preceded the refinement in Refmac due to the lower resolution of the map. MDFF calculations were performed as described in the tutorial. No refinement of molecular dynamics simulations was performed for complex with antigen D.

## Conclusions

Delivering high quality cryo-EM structures for small sized complexes at sub-100 kDa range requires an optimized workflow with strategic decision making, especially when preferred orientation occurs. Here, we shared a few representative cases to demonstrate how cryo-EM can be routinely applied to reveal details of epitope-paratope interface quickly and reliably and support broad project needs. Coupled with strong computational structure modeling, we were able to extract accurate structural information using this workflow from less optimized cryo-EM maps within a short time. The workflow turns out to be much more effective compared to spending time and resources to improve resolution. In our work, we tried to overcome preferred orientation by applying commonly used techniques developed for large sized targets, however, for particles in the sub-100 kDa size range, we observed that preferred orientation has to be resolved through new complex formation, such as adding additional Fab, V_HH_, or another binding partner to a complex as shown in the case study for antigen B. In extremely challenging cases, knowledge of glycosylation and protein features of antigens can provide critical guidance to get meaningful structural information when high resolution 3D volume reconstruction is not feasible.

## Acknowledgments

We want to thank cryo-electron microscopy facilities of University of Massachusetts, Harvard Medical School, University of Michigan for their help with sample screening and data acquisition; and Robert Cost for proofreading our draft and providing valuable comments.

